# Reproductive neuronal circuitry in adaptive changes of energy balance

**DOI:** 10.1101/2021.09.09.459635

**Authors:** Pilhwa Lee, Cristina Sáenz de Miera, Nicole Bellefontaine, Marina A Silveira, Thais T Zampieri, Jose Donato, Kevin W. Williams, Renata Frazao, Carol F. Elias

**Author notes:** Corresponding Author: Carol F. Elias, Ph.D. 1137 E Catherine St. 7725 Med Sci II, Ann Arbor, MI 48109. first co-authors (Pilhwa Lee, Cristina Sáenz de Miera).

## Abstract

The crosstalk between metabolism and reproduction is essential for species survival. When dysfunctional, this interaction may decrease reproductive efficiency, but in physiological conditions of high energy demands, e.g., pregnancy and lactation, it is highly beneficial. Females display adaptive responses that assure offspring survival and health, including increased food intake and suppression of the reproductive function. Some of these physiological responses are modulated by leptin actions in neuronal pathways that are still unclear. The hypothalamic ventral premammillary nucleus (PMv) is a key integrative node of metabolic cues and reproductive status, comprised of either leptin-depolarized or -hyperpolarized neurons. Here, we show that the subset of leptin-hyperpolarized neurons coexpresses dopamine transporter (DAT) and prolactin receptor. DAT expression is higher in prepubertal conditions, when reproductive function is suppressed. These neurons innervate AgRP presynaptic terminals and may potentiate their inhibitory actions on reproduction. We further applied a mathematical model to reconcile our new findings with the current literature and to verify if those neurons are putative components of the metabolic control of reproduction. In our model, leptin-depolarized PMv neurons project to and directly stimulate kisspeptin and gonadotropin releasing hormone (GnRH) neurons. Leptin-hyperpolarized PMv DAT neurons are directly stimulated by prolactin and project to inhibitory control sites. During conditions of high prolactin levels, i.e., late pregnancy and lactation, this pathway may overcome the former, facilitating AgRP actions in the suppression of the reproductive function. Our model also predicts that overstimulation of this pathway may underlie earlier puberty and reproductive deficits observed in conditions of metabolic dysfunction.

**Significance Statement:** Women with excess or low energy stores (e.g., obesity or anorexia) have reproductive deficits, including altered puberty onset, disruption of reproductive cycles and decreased fertility. If able to conceive, they show higher risks of miscarriages and preterm birth. The hypothalamic circuitry controlling the interplay between metabolism and reproduction is undefined. Neurons in the ventral premammillary nucleus express leptin receptor and project to reproductive control sites. Those neurons are essentially glutamatergic, but functionally and phenotypically heterogeneous. They either depolarize or hyperpolarize in response to leptin. We show that leptin-hyperpolarized neurons coexpress dopamine transporter and prolactin receptor, and project to AgRP inhibitory output. Computational modeling was applied to build a neuronal network integrating metabolism and reproduction in typical and dysfunctional physiology.

## Introduction

Pubertal development and the maintenance of the reproductive function are disrupted in states of negative energy balance or excess energy reserve (1-3). If energy stores are low, puberty is delayed, the reproductive cycles are prolonged, and sub- or infertility ensue (4-6). High adiposity, on the other end, induces earlier pubertal development and decreased fertility in adult life (7-12). Although apparently disruptive, the crosstalk between metabolism and reproduction is highly beneficial in physiological conditions of high energy demands. For example, during pregnancy and lactation, an adaptive strategy that assures the nutritional needs for offspring survival and health has emerged. Increased caloric intake and temporary suppression of the reproductive function are orchestrated by a complex neuronal network modulated by circulating hormones and metabolic cues (13, 14). Among them, leptin and prolactin have critical roles (15, 16). Leptin signaling-deficient subjects develop obesity and remain in an infertile prepubertal state (17-21). In mice, direct leptin actions only in the brain are sufficient to normalize body weight, to induce puberty and maintain fertility (22-25). Chronic prolactin administration disrupts the reproductive axis and increases body weight (15, 26, 27). In physiological conditions of high energy demands, increasing levels of prolactin and hypothalamic leptin resistance allow for energy accumulation and adequate nutrition of the fetus and newborn (13, 16, 26, 28).

The ventral premammillary nucleus (PMv) expresses a dense collection of leptin receptor (LepRb) neurons, and is recognized as an important hypothalamic site in the metabolic control of the reproductive function (24, 29-32). Bilateral lesions of the PMv disrupt estrous cycles and the ability of leptin to increase luteinizing hormone secretion following a fast (5). Endogenous restoration of LepRb exclusively in PMv neurons rescues pubertal maturation and fertility in LepRb *null* female mice (24). The PMv LepRb neurons, however, do not comprise a homogeneous population, i.e., about 75% depolarize and 25% hyperpolarize in response to leptin (33), but their seemingly dissociated function is poorly understood.

The PMv neurons are essentially glutamatergic and innervate brain sites associated with reproductive control, sending direct inputs to kisspeptin and gonadotropin releasing hormone (GnRH) neurons (24, 30-32, 34). A subset of PMv neurons also expresses the dopamine transporter (DAT), a membrane protein associated with dopamine reuptake at the presynaptic terminals (35-37). PMv DAT neurons are unique in the sense that they do not release dopamine; rather, they are essentially glutamatergic (36, 38). Remote manipulation of PMv DAT neuronal activity has shown an action in male social behavior and inter-male aggression (36, 37). However, the role of PMv DAT neurons in female physiology or motivated behaviors have not been described, and whether they coexpress LepRb and/or have a function in the metabolic control of the reproductive function is unknown.

In this study, we show that DAT is expressed in the subpopulation of LepRb neurons hyperpolarized by leptin that also coexpresses prolactin receptor. We further applied a mathematical model to verify if those neurons are putative components of the hypothalamic circuitry associated with the metabolic control of pubertal development and with the suppression of the reproductive neuroendocrine axis typical of pregnancy and lactation, when prolactin levels are high. Our model predicts that overstimulation of this pathway underlies the reproductive restraint observed in adaptive physiology (e.g., pregnancy and lactation) and in conditions of metabolic dysfunction.

## Results and Discussion

### DAT mRNA expression in the PMv is higher in prepubertal mice

The *Slc6a3* (DAT) gene is expressed in the PMv of male and female mice (35, 36). To assess potential sexual dimorphism, we generated the DAT-Cre tdTom mice (39). No difference in the number of cells was observed in male *vs* female PMv (*p* = 0.09, Figure 1A-B). However, PMv DAT mRNA was higher in diestrous female compared to male mice (Figure 1C), suggesting that developmental processes drive DAT expression in PMv neurons which is masked by the Cre-induced reporter gene.

**Figure 1.**
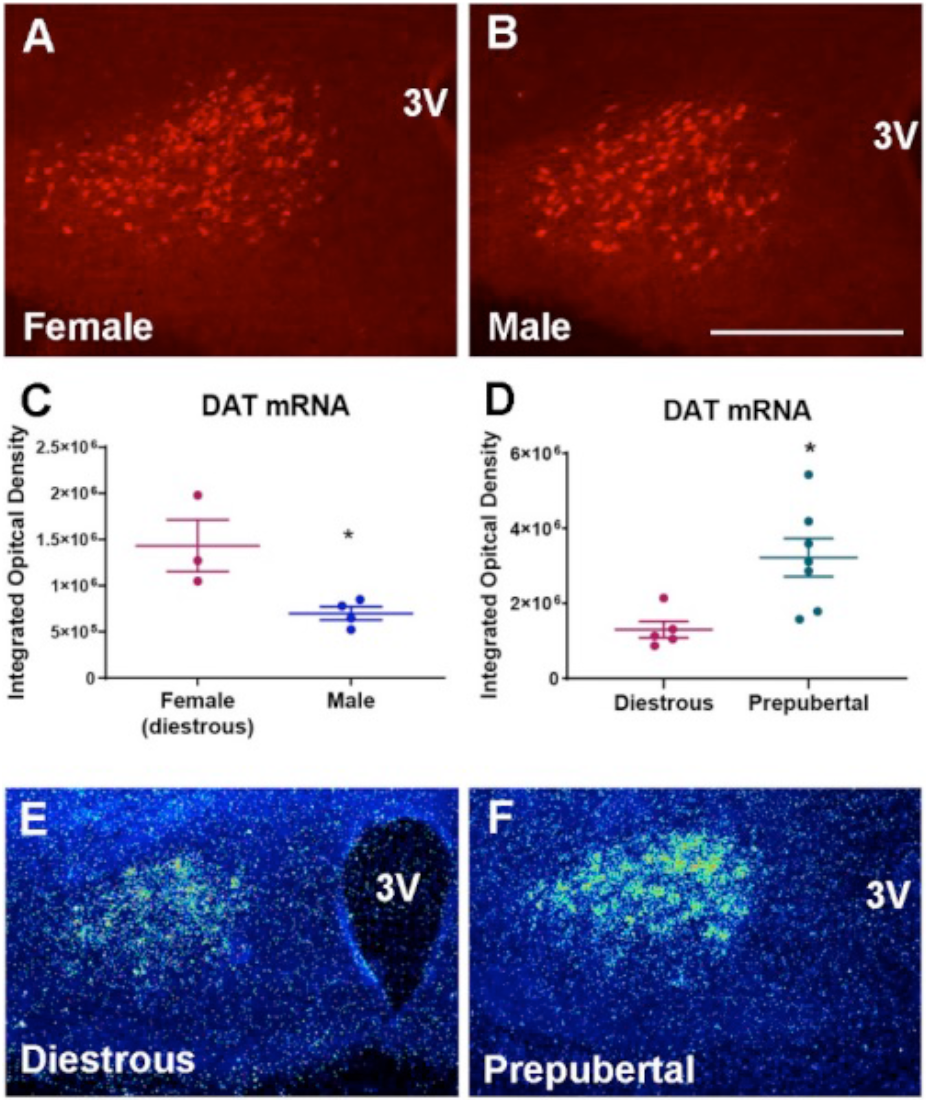
PMv *Slc6a3* (DAT) gene expression varies with sex and development. A-B. Fluorescent image showing DAT-Cre tdTomato cells in the PMv of female (A) and male (B) adult mice. C-D. Graphs showing the quantification of the DAT hybridization signal (silver grains) in the PMv by integrated optical density (IOD). E-F. Pseudocolor of dark field images showing intensity variation in DAT mRNA expression. Unpaired t-test, *p* < 0.05. Scale bars: A-B = 200 µm, E-F = 300 µm.

Prepubertal females (postnatal day 19, P19) have higher DAT mRNA expression compared to diestrous mice (P60-70, Fig. 1D-H). To assess if the reduction of PMv DAT mRNA in adult mice is a result of increasing estradiol (E2) levels, hypothalamic sections from diestrous, ovariectomized (OVX), and OVX + E2 mice were analyzed. We found no differences between the 3 groups (n=4 mice/group, one-way ANOVA, *p* = 0.54), indicating that DAT mRNA is responsive to factors other than E2 during pubertal maturation.

### A subset of PMv DAT neurons is responsive to leptin

Leptin action only in PMv neurons is sufficient to induce puberty and improve fertility in LepRb-*null* mice (24). To assess if PMv DAT neurons are direct targets of leptin, DAT-Cre tdTom mice received an intraperitoneal (ip.) injection of leptin, and immunoreactivity for phosphorylation of signal transducer and activator of transcription 3 (pSTAT3-ir), an intracellular signal directly triggered by leptin (40), in tdTom neurons was quantified. We found that 32.5 ± 3.9% tdTom positive cells in female and 44.2 ± 2.1% in male mice coexpress pSTAT3-ir (Figure 2A-B).

**Figure 2.**
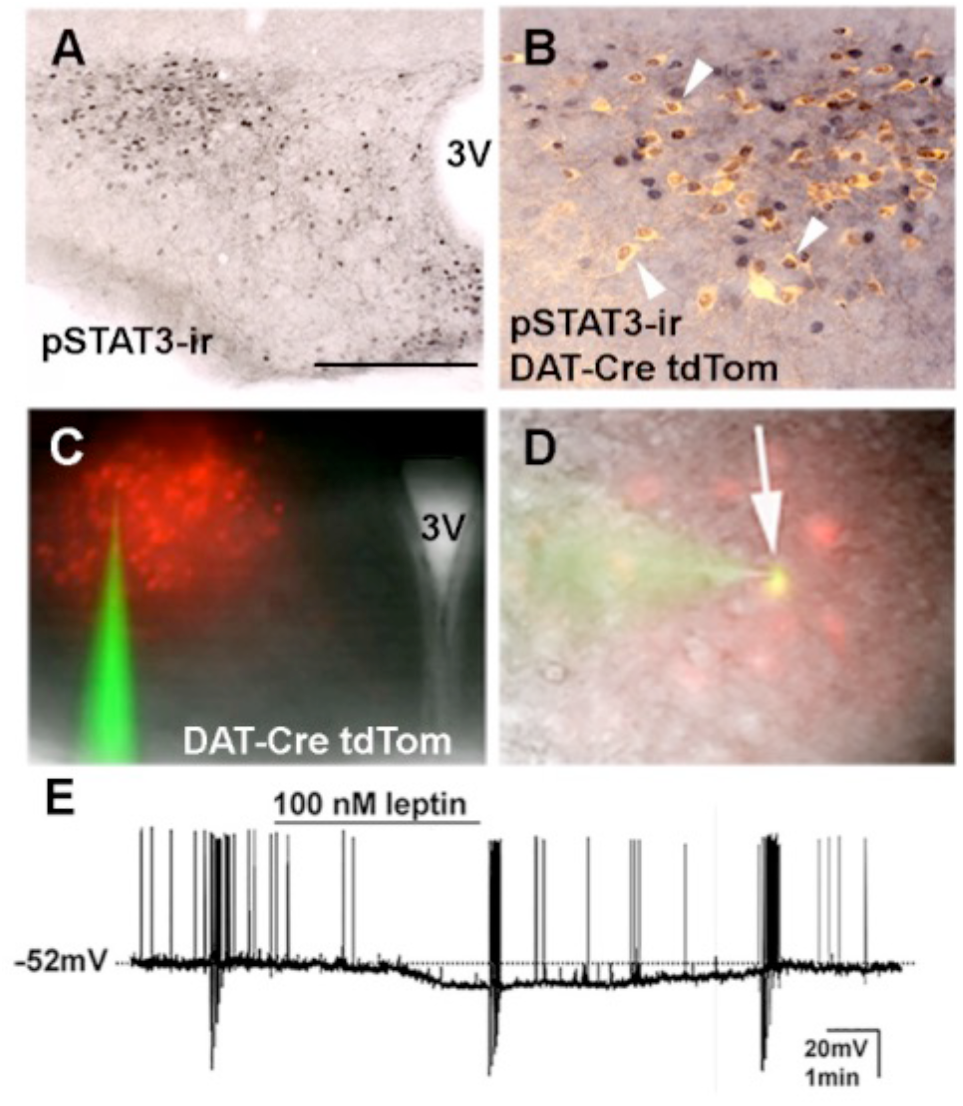
A subset of PMv DAT neurons is responsive to leptin. (A) Bright field image showing the distribution of leptin-induced pSTAT3-ir in the PMv. (B) High magnification of merged fluorescent and brightfield images showing the colocalization of phosphorylation of STAT3 immunoreactivity (pSTAT3-ir, black nuclei) and tdTomato positive cells (pseudocolor yellow cytoplasm) in the PMv of a female mouse. (C) Fluorescent image showing the PMv in the brain slices, recognized by tdTomato expression in DAT-Cre neurons. (D) Merged image showing the colocalization between a recorded DAT-Cre tdTomato neuron (red) and the AF488 dye (green), dialyzed during the recording. (E) Representative current-clamp recording demonstrating leptin (100 nM) induced hyperpolarization in a subset of DAT-Cre tdTomato neurons. The dashed line indicates resting membrane potential (−52 mV). Scale bars: A, C = 400 µm, B = 200 µm, D = 50 µm.

To investigate the effect of leptin on the membrane excitability of DAT neurons, we performed current clamp recordings of tdTom neurons. The average resting membrane potential (RMP) of recorded neurons was −52.8 ± 2.0 mV (range from −62 to −38 mV, 16 cells out of 8 male adult mice). We found that bath application of leptin hyperpolarized about 30% of the recorded neurons (4 out of 14 cells) whereas the RMP of 70% of the recorded cells remained unchanged in response to leptin (Figure 2C-E). The leptin-hyperpolarized cells in PMv exhibited −7.8 ± 0.8 mV change in the RMP by comparing baseline V_rest_ to treatment V_rest_ (from −56.2 ± 2.1 mV to −64.0 ± 2.1 mV; two-tailed *t*-test, *p* < 0.002, Figure 2E). The hyperpolarization was accompanied by a 20% decrease in whole-cell input resistance, from 811.8 ± 139.5 MΩ during baseline to 642.3 ± 110.1 MΩ following leptin application (*p* = 0.01). The similarity between the percentage of DAT-Cre neurons expressing leptin-induced pSTAT3-ir and of those hyperpolarized by leptin is striking. Although neurons may be responsive to leptin in ways we have not tested in this study, our findings indicate that leptin action in PMv DAT neurons is mostly inhibitory.

The higher expression of DAT transcript in the PMv of prepubertal mice, the role of PMv neurons in reproduction, and the hyperpolarizing effect of bath application of leptin on PMv DAT-Cre neurons suggested that PMv DAT/LepRb neurons are more active in conditions of reduced activity of the reproductive axis. Among them, pregnancy and lactation are physiological states of reproductive restraint, also characterized by high circulating levels of prolactin (41-44).

### DAT, LepRb and prolactin receptor (PrlR) are coexpressed in PMv neurons

A subset of PMv LepRb neurons are responsive to prolactin (45), but whether they coexpress DAT has not been determined. Because DAT mRNA expression is higher in prepubertal mice, Cre-induced reporter gene is not an adequate marker for DAT expression in adult mice. Fluorescent *in situ* hybridization was used to assess transcript coexpression in adult females on diestrus. About half of PMv LepR neurons coexpressed DAT mRNA (58.6 % ± 4.1), whereas virtually all DAT neurons coexpressed LepR mRNA (93.1 % ± 1.9, Fig 3A-B). About a quarter of LepR (26.2 ± 3.5%) and of DAT (25.5 ± 1.3%) neurons coexpressed PrlR mRNA (Fig 3A-D). Neurons expressing only LepR or PrlR were also observed. Because most of PMv DAT neurons coexpressed LepR in adult females, our findings indicate that prolactin action in PMv LepRb neurons preferentially target DAT expressing cells, the subpopulation of neurons hyperpolarized by leptin.

**Figure 3.**
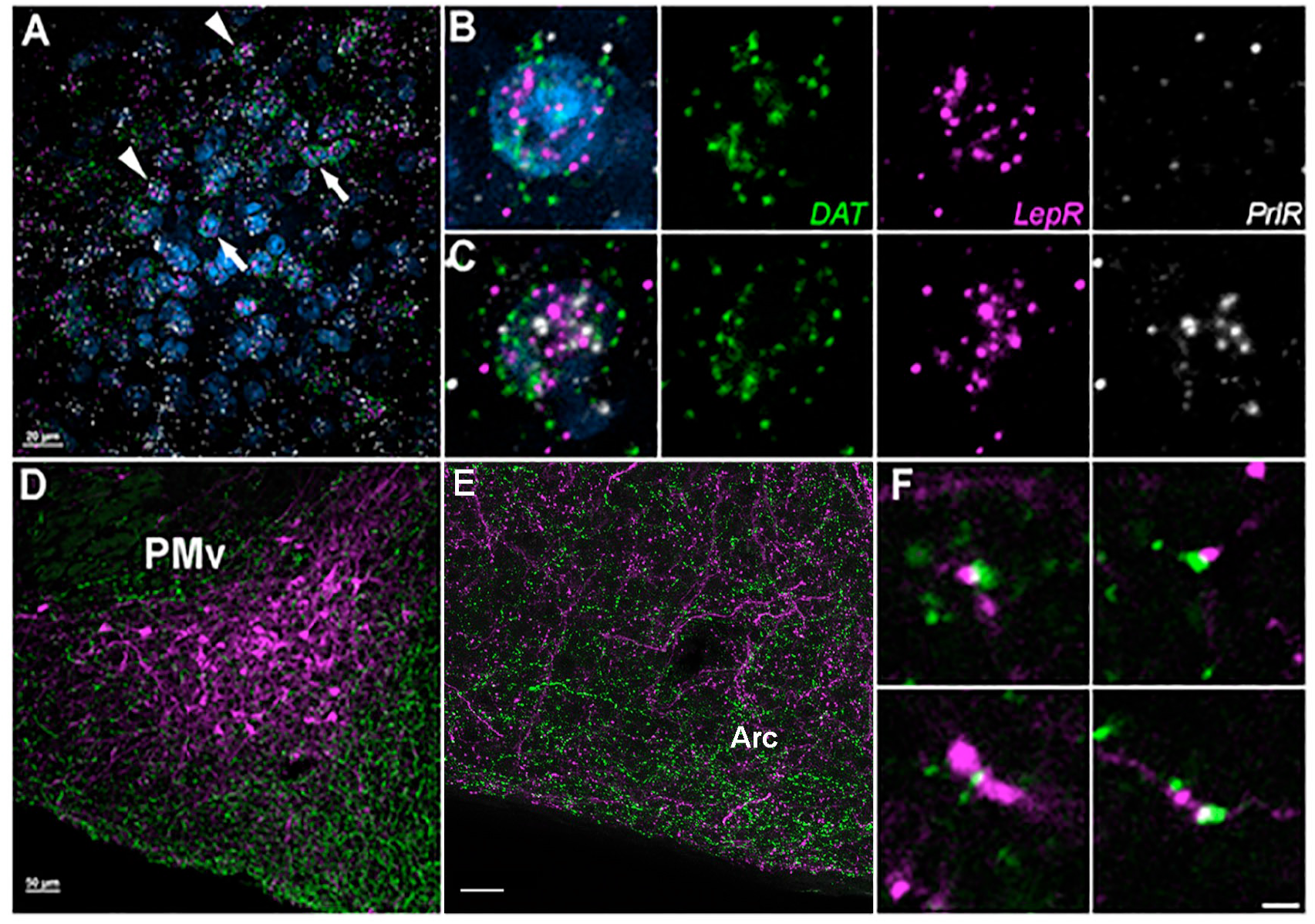
LepR, DAT and PrlR are coexpressed in PMv neurons. (A) Fluorescent image showing representative high magnification fluorescent in situ hybridization image depicting the colocalization of LepR (magenta), DAT (green) and PrlR (white) in the PMv of a diestrous female. Arrowheads point to cells co-expressing DAT, LepR and PrlR mRNA. Arrows point to cells co-expressing LepR and DAT mRNA. Blue = DAPI. (B-C) Higher magnification of an individual cell from the region depicted in A, co-expressing DAT and LepR mRNA, but with no PrlR mRNA (B), or all three transcripts (C). Left: merged channels image. Individual channels shown in the right panels. (D-F) PMv DAT projections are in close contact with AgRP fibers. (D) Fluorescent image showing AAV synaptophysin mCherry injection site centered in the PMv (magenta) and AgRP immunoreactive fibers (green) in an adult DAT-Cre female mouse. (E) High magnification 10 µm thick Z-stack image at the posterior and lateral level of arcuate nucleus (Arc), anterior to the injection site showing close appositions between mCherry- and AgRP-immunoreactive fibers. (F) High magnification showing examples of close contacts between mCherry- and AgRP-immunoreactive fibers in a single Z plane (0.75-µm thick) in the region depicted in E. Scale bars: A, E = 20 µm, D = 50 µm, F = 2 µm.

### PMv DAT cells project to AgRP neuronal target sites

We and others have shown that PMv neurons are essentially glutamatergic and innervate GnRH and Kiss1 neurons (24, 31, 32, 34). Due to the hyperpolarizing effect of leptin on PMv DAT neurons, we hypothesized that this neuronal population innervates neurons associated with inhibition of the neuroendocrine reproductive axis. A Cre-dependent synaptophysin mCherry AAV was delivered unilaterally into the PMv of DAT-Cre female mice (Fig. 3E). Strong projections to the lateral arcuate nucleus (Arc) and the VMHvl were observed. We then assessed if PMv DAT terminals innervate AgRP neurons, a key component of leptin physiology and an inhibitory player in the metabolic control of reproduction (25, 46, 47). *Agrp* expression is high in negative energy balance, and remote activation of AgRP increases food intake and inhibits the reproductive function (46-49). Close appositions were mostly observed between mCherry- and AgRP-immunoreactive fibers in the lateral Arc and in the ventromedial aspects of the VMHvl (Fig. 3F). Little or virtually no innervation of AgRP cell bodies was observed.

### PMv integration of energy balance and reproduction

Based on published literature and our current findings, we built a model to gain insights into the role of PMv neuronal circuitry in the integration of energy balance and reproductive function (Fig. 4A). The PMv LepRb neurons are essentially glutamatergic and comprise 2 distinct populations: one depolarized (excited) and one hyperpolarized (inhibited) by leptin. Leptin-excited PMv neurons project to Kiss1 and GnRH neurons, inducing their activation via glutamatergic neurotransmission (24, 30-32). Kiss1 neurons are crucial for pubertal development and fertility with a major role in GnRH secretion (21, 50-52). Leptin-inhibited PMv DAT neurons, on the other hand, innervate AgRP fibers that synapse onto and inhibit Kiss1 neurons via GABAergic neurotransmission (46). The direct projections from leptin-inhibited PMv-LepRb/DAT neurons to Kiss1 and/or GnRH neurons are assumed to be weak.

**Figure 4.**
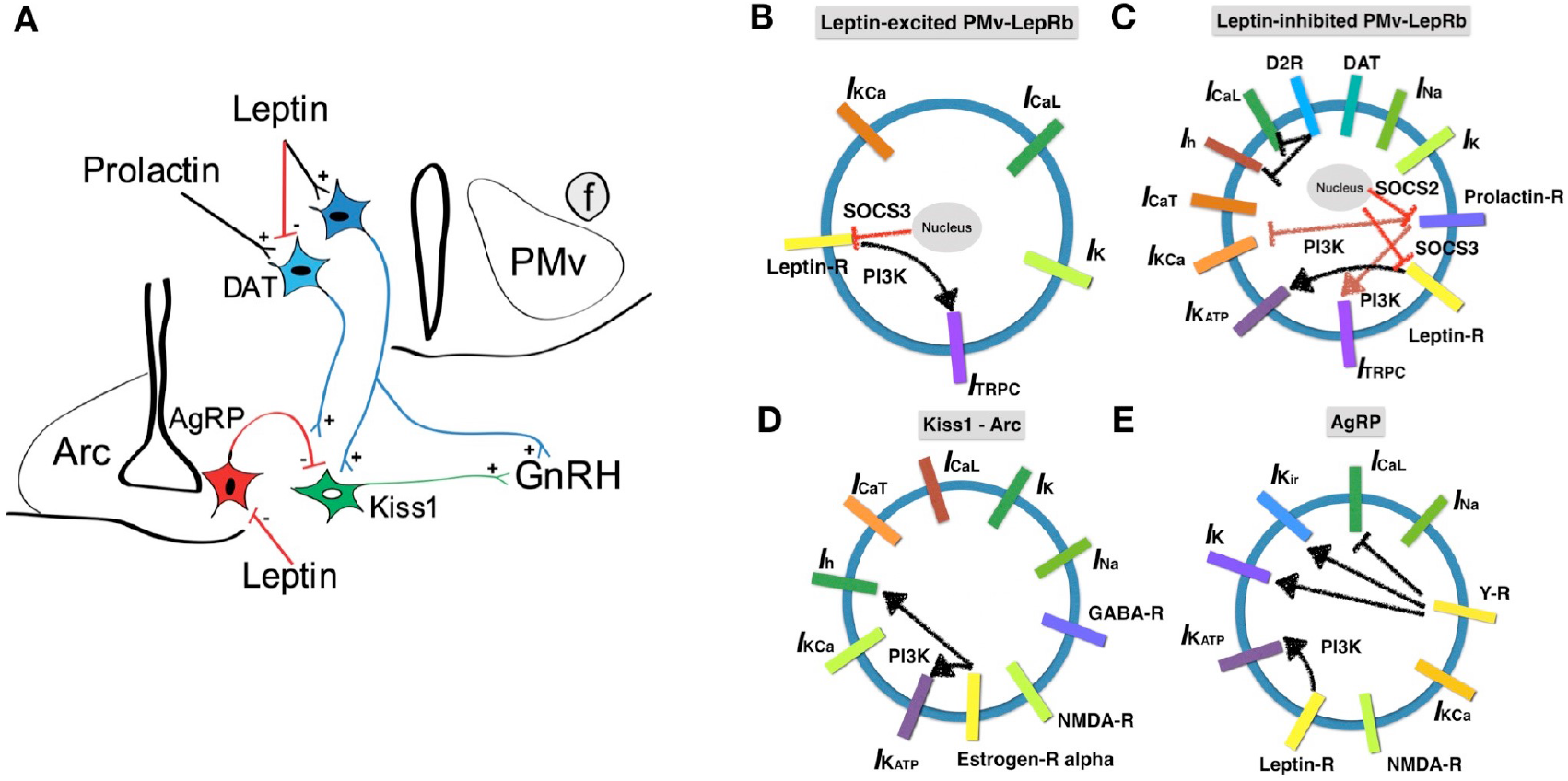
Proposed model for PMv neuronal circuitry in the metabolic control of the reproductive function. (A) Upstream input signals from leptin and prolactin hormones. Subpopulations of PMv-LepRb differentially respond to acute leptin, one depolarizes (excites) and the other hyperpolarizes (inhibits). Leptin-excited PMv-LepRb neurons innervate GnRH neurons directly and indirectly via Kiss1 neurons. Leptin-inhibited LepRb neurons coexpress DAT and PrlR, and project to AgRP fibers that inhibit Kiss1 neurons. (B-E) Ionic conductance and hormone receptors. In leptin-excited PMv-LepRb neurons (B), TRPC channels are activated by leptin via PI3K pathway. In leptin-inhibited PMv-LepRb/DAT neurons (C), K_ATP_ channels are activated by leptin and TRPC channels are activated by prolactin via PI3K pathway. Also included is the suppression of LepR by SOCS3 and of PrlR by SOCS2. In AgRP and Kiss1 neurons (D-E), the estrogen receptor-α (ERα) and NPY/AgRP receptors are also illustrated in their connection with ionic conductance, but the actual signal sensitivities are beyond the scope of this article.

The depolarizing effects of prolactin on PMv neurons (53), the prolactin-induced phosphorylation of STAT5 in PMv-LepRb neurons (45) and the colocalization of DAT and PrlR mRNA in a subset of PMv LepR neuronal population led us to postulate that PMv-LepRb/DAT neurons exert a key role in adaptive physiology, when suppression of the reproductive axis and high prolactin levels are apparent, e.g., during pregnancy and lactation (Fig. 4A).

### Modeling conductance-based electrophysiology

We have developed a conductance-based electrophysiology model for 4 types of neuron: 1) Leptin-excited PMv-LepRb neurons; 2) Leptin-inhibited PMv-LepRb neurons; 3) Kiss1 neurons in the arcuate nucleus; and 4) AgRP neurons. The population-level activity in the same type of neuron was treated as homogeneous and scaled in the presynaptic strength 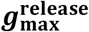 (i.e., presynaptic glutamatergic maximum release to Kiss1) for the synaptic transmission from one type to another (Equation [1]). For each type of neuron, ion channels and intracellular signaling are illustrated in Fig. 4B-E. The presynaptic release of glutamate is formulated based on the energetics of exocytosis events in terms of membrane voltage:

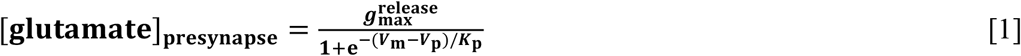

where ***V***_**p**_ is the threshold membrane potential for release of glutamate, and ***K***_**p**_ is the spreading factor in the sigmoidal response (54, 55). The kinetic release of GABA neurotransmitter is prescribed similarly.

In postsynaptic neurons, glutamatergic NMDA receptors are gated with binding of glutamate, and formulated as follows:

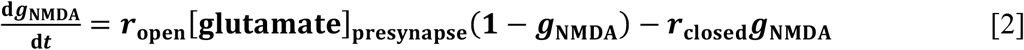

Nearly all the current elicited by glutamatergic EPSCs in Kiss1 and AgRP neurons is mediated by NMDA receptors, with only a minor contribution from AMPA receptors (56, 57). Therefore, in the current model with minimal components, we considered only NMDA receptors in Kiss1 and AgRP neurons for glutamatergic innervation, and we modelled NMDA receptors with a non-Mg dependent conductance (i.e. without Mg^2+^ block). AMPA receptors could be added in the future to further fine-tune the model. The kinetics of GABA receptors are treated similarly (51, 52).

Upon leptin and prolactin binding, the expression of suppressor of cytokine signaling 3 (SOCS3) and 2 (SOCS2) are induced, respectively, exerting negative feedback regulation on receptor signaling (Fig. 4B, C) (58, 59, 60, 61). The effective impact of leptin and SOCS3, and of prolactin and SOCS2 were modeled by competitive binding shown in Equations [5], [25] and [39] in Supplemental Information (SI Appendix).

#### 1) Leptin-depolarized PMv-LepRb neurons

We have prescribed minimal components of ionic currents to generate repetitive spiking action potentials as follows:

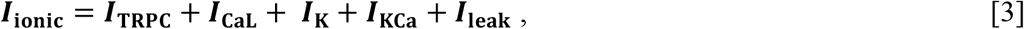

where the fast dynamics of membrane voltage is coupled to the slow dynamics of calcium handling in endoplasmic reticulum as well as *I*_KCa_ (calcium-activated potassium currents) to generate oscillatory spiking profiles. *I*_TRPC_ is determined by leptin-induced PI3K signaling and feedbacks via SOCS3 (33, 61, 62) (Fig. 4B, SI Appendix Equation [5] and [6]). Leptin-depolarized PMv-LepRb neurons project to Kiss1 neurons releasing glutamate. The 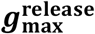 is prescribed to be proportional to the PMv-LepRb subpopulation size, where the neurons in the subpopulations are assumed to be identically stimulated.

#### 2) Leptin-hyperpolarized PMv-LepRb neurons

For leptin-hyperpolarized PMv LepRb/DAT neurons, we have prescribed the following ionic currents:

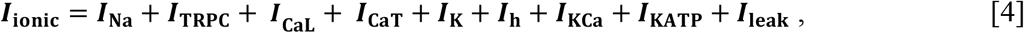

where *I*_KATP_ (ATP-dependent potassium currents) is also determined by leptin via PI3K signaling and the feedback by SOCS3 (Fig. 4C, SI Appendix Equation [37]). *I*_KCa_ is affected by prolactin (62) and SOCS2 (SI Appendix Equation [36]). Dopamine uptake by DAT effectively reduces dopamine availability and binding to dopamine D2 receptor (D2R) (63). As D2R inhibits *I*_CaL_ (L-type calcium current) and *I*_h_ (hyperpolarization-activated current) (64), DAT would facilitate the activation of *I*_CaL_ and *I*_h_ currents transmitting glutamatergic signals to AgRP neurons. Upon membrane hyperpolarization, an inhibitory rebound is invoked by *I*_h_ and *I*_CaT_ (low-threshold T-type calcium current) currents (37, 54).

#### 3) Kiss1 neurons

Kiss1 neurons in the Arc are directly innervated by AgRP fibers. Optogenetic stimulation of AgRP neurons evoked IPSCs in recorded Kiss1 neurons which were blocked by a GABA_A_ receptor antagonist (46). In the absence of AgRP/GABA innervation, Arc Kiss1 neurons display lower frequency of miniature inhibitory postsynaptic currents (mIPSC), thereby decreasing inhibitory inputs (46). Thus, according to our model, activation of AgRP fibers via increased glutamatergic neurotransmission from LepRb/DAT neurons increases the inhibitory input to Arc Kiss1 neurons (Fig 4A). The ionic currents are composed of the following components:

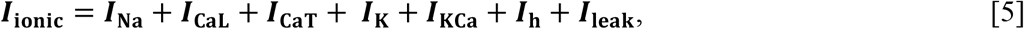

where *I*_h_ and *I*_CaT_ currents also play a role (65). In the proposed neuronal circuitry, leptin-inhibited PMv-LepRb/DAT innervation of AgRP fibers functions as a presynaptic modulator of Arc Kiss1 neuronal activity.

#### 4) AgRP neurons

AgRP neurons are directly inhibited by leptin (66). Of note, leptin hyperpolarization of both PMv LepRb and AgRP neurons requires the activation of K_ATP_ channel via PI3K signaling (33, 66, 67). The PMv-LepRb/DAT (glutamatergic) terminals are in close apposition with AgRP fibers. Therefore, depolarization of PMv-LepRb/DAT neurons may induce AgRP and/or GABA release from presynaptic terminals innervating Arc Kiss1 neurons. The ionic currents involved include:

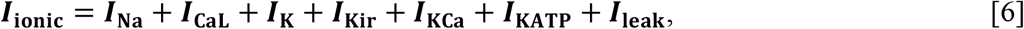

where *I*_KATP_ is determined by leptin and SOCS3 (SI Appendix Equation [79]). In physiological conditions of reproductive restraint and high prolactin levels, suppression of Kiss1 neurons (68, 69) may be potentiated by decreased inhibition of AgRP neurons by leptin (resistance) and activation of leptin-inhibited PMv-LepRb/DAT neurons by prolactin.

### Plasticity of leptin-excited and leptin-inhibited PMv neurons in pubertal development

The difference between the number of DAT-Cre tdTom neurons and DAT mRNA expression in adult mice suggests PMv-LepRb neurons display developmental plasticity associated with the metabolic control of sexual maturation. To model energy sensing in the PMv neuronal circuitry, we reconstructed the bursting and spiking profiles prescribing 100nM of leptin and 100nM of SOCS3 for both leptin-activated and leptin-inhibited PMv neurons (Fig. 5A-B). The membrane potential is more hyperpolarized, but bursting activity from hyperpolarization-activated *I*_h_ current and low-threshold T-type calcium channel (*I*_CaT_) is engaged. The downstream inputs to Kiss1 and AgRP/GABA were reconstructed focused on their spiking and bursting membrane voltages. Due to the developmental plasticity of PMv-LepRb/DAT neurons we assumed the proportion of excited/inhibited neurons are modified during pubertal transition. In adult females, 75% of PMv-LepRb neurons are depolarized and 25% are hyperpolarized by leptin (33). In prepubertal mice, according to data from DAT-Cre tdTom *vs* DAT mRNA we assumed 50% of PMv-LepRb neurons are either depolarized (non-DAT) or hyperpolarized (DAT) by leptin. Under these conditions, AgRP neurons are more depolarized compared to adults, when they constitute 25% of the total LepRb expressing neurons in the PMv (adult, Fig 6A-B). When above −20 mV, the presynaptic GABA release threshold (up and down) (55), AgRP neurons increase IPSCs in Kiss1 neurons. As a result, the spiking activities of Kiss1 neurons are suppressed when leptin-inhibited neurons encompass 50% compared to 25% of PMv-LepRb/DAT neurons in adults (Fig. 6D-E). According to our model, at least part of the leptin actions in pubertal development is due to the reduction in the proportion of PMv-LepRb/DAT neurons that are inhibited by leptin (from 50% to 25%). During pubertal transition, this change decreases the modulatory strength exerted upon AgRP/GABA innervation of Kiss1 neurons, releasing the latter from activity restraint. If the PMv-LepRb/DAT neuronal plasticity during pubertal transition is compromised, we predict AgRP/GABA inhibitory actions on Kiss1 neurons will persist and the Kiss1 neuronal firing rate will remain low, leading to delayed puberty. This mechanism may add to the pubertal delay observed in states of AgRP upregulation (70).

**Figure 5.**
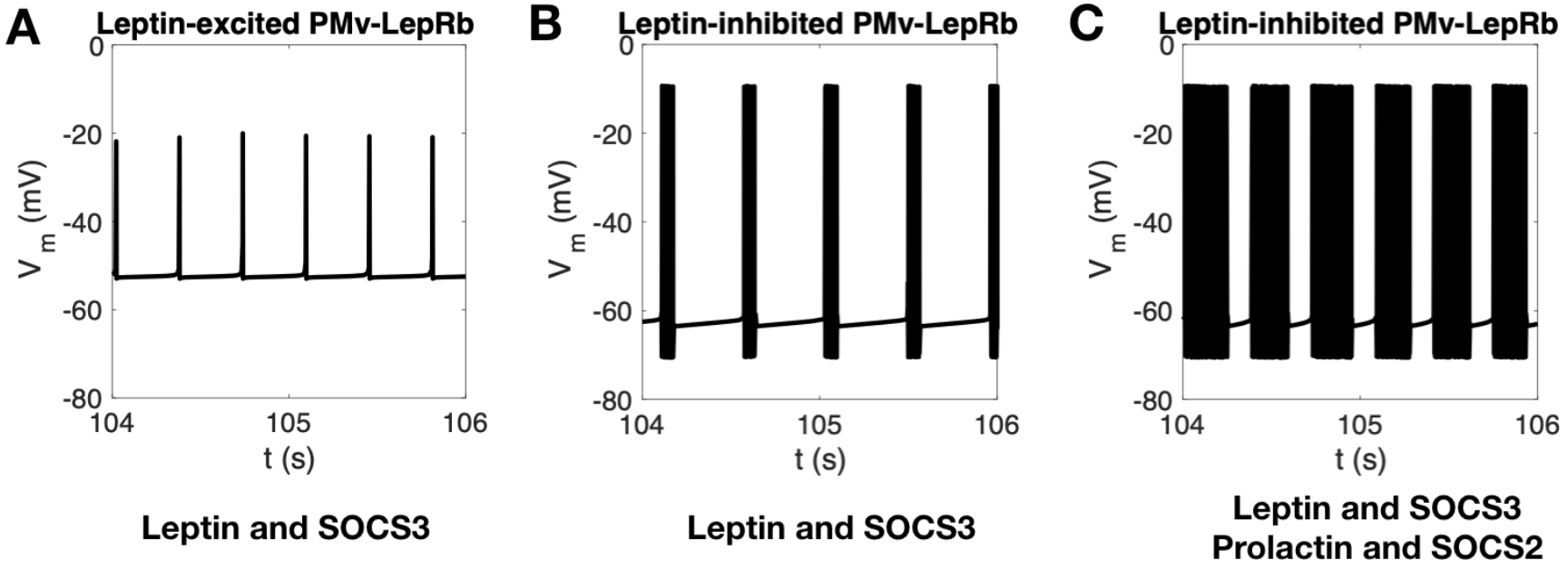
Bursting and spiking profile of PMv neurons. (A) The spiking profile of leptin-depolarized PMv-LepRb neurons with 100 nM of leptin and 100 nM SOCS3. (B) The bursting profile of leptin-hyperpolarized PMv-LepRb neurons with 100 nM of leptin and 100 nM of SOCS3. The membrane potential baseline is lowered with hyperpolarization, but there are bursting activities from hyperpolarization-activated (*I*_h_) channels and low-threshold T-type (*I*_CaT_) calcium channels. (C) The spiking profile of leptin-hyperpolarized PMv-LepRb neurons with 100 nM of leptin, 100 nM of SOCS3, 5 μM of prolactin, and 1.0 μM of SOCS2.

**Figure 6.**
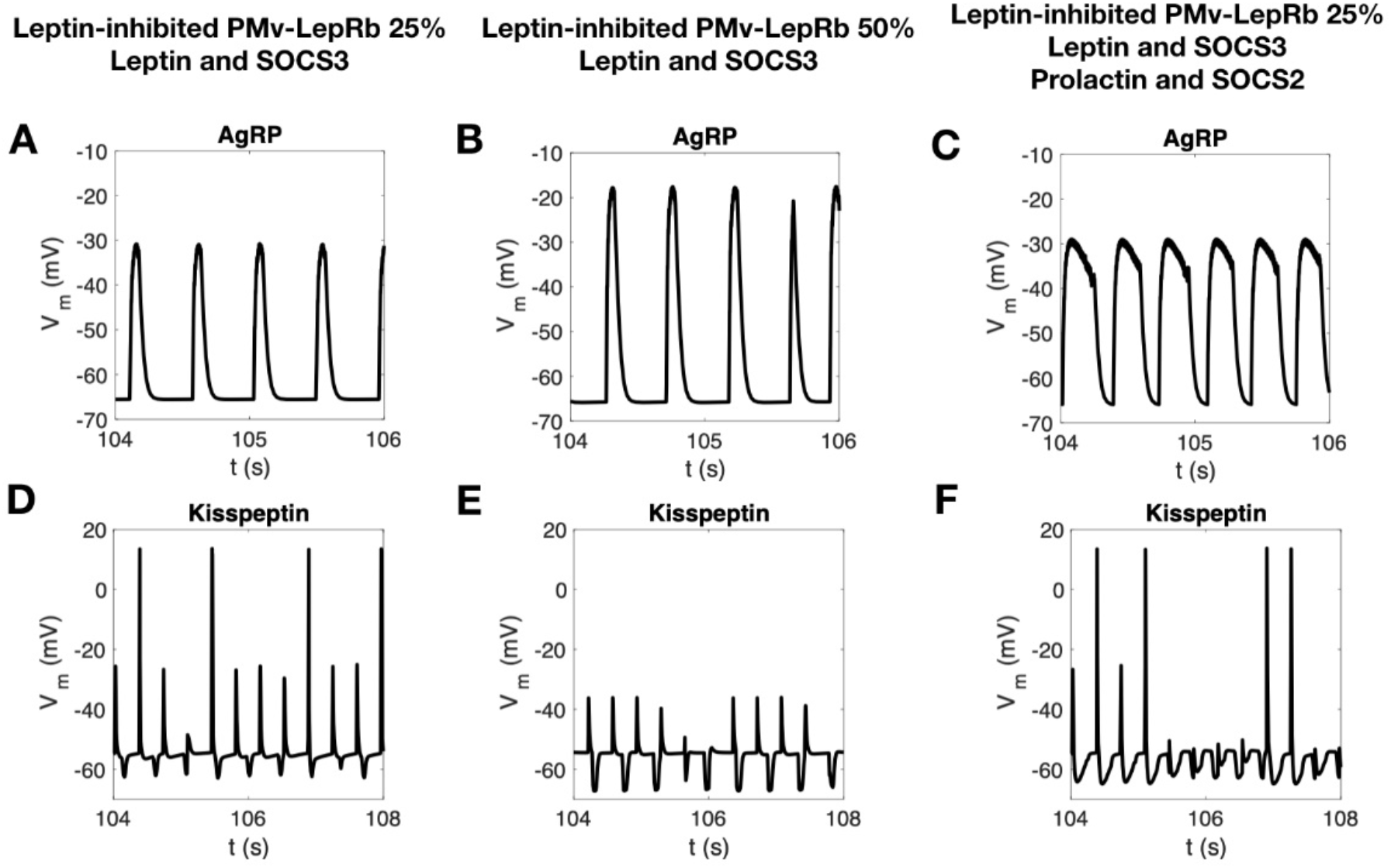
Bursting and spiking profiles of downstream Kiss1 and AgRP neurons. (A, D) Representation of bursting and spiking profile in adult female mice with a 75% leptin-excited and 25% leptin-hyperpolarized PMv-LepRb neurons balance using 100 nM of leptin and 100 nM of SOCS3. (B, E) Representation of bursting and spiking profile in prepubertal stage with a 50% leptin-depolarized and 50% leptin-hyperpolarized PMv-LepRb neurons balance, using 100 nM of leptin 100 nM of SOCS3. (C, F) Representation of bursting and spiking profile in pregnancy and lactation, with a 75% leptin-hyperpolarized and 25% leptin-hyperpolarized PMv-LepRb neurons balance with 100 nM of leptin, 100 nM of SOCS3 prescribed for leptin-depolarized PMv-LepRb neurons and 100 nM of leptin, 100 nM of SOCS3, 5 μM of prolactin, and 1.0 μM of SOCS2 prescribed for leptin-hyperpolarized PMv-LepRb neurons.

While recapitulating the pubertal development, we did not include the progressive change in leptin responsiveness and K_ATP_ channel expression in AgRP neurons from depolarizing (40% at P15) to hyperpolarizing (66% at P30) (71) because it is not possible to determine the neuronal origin and phenotype of AgRP presynaptic terminals innervated by DAT fibers. Our model was focused on the roles of population-level plasticity in leptin-depolarized and -hyperpolarized PMv-LepRb neurons.

### PMv-LepRb neuronal response to changing levels of circulating prolactin

During late pregnancy and lactation, prolactin secretion by the pituitary gland is increased and hypothalamic SOCS2 levels are elevated up to 10-fold (16, 72). PrlR and LepRb are co-expressed in hypothalamic nuclei of adult female mice (45). Particularly for the PMv, about 25% of leptin-inhibited neurons coexpress PrlR in adult females.

We have modeled PMv LepRb neuronal circuitry during late pregnancy and lactation by prescribing 100 nM of leptin, 5.0 μM of prolactin, 100 nM of SOCS3, and 1.0 μM of SOCS2 (72). Based on the predicted low expression of PrlR in leptin-depolarized PMv-LepRb neurons, leptin and prolactin effects are only applied to leptin-hyperpolarized PMv-LepRb/DAT neurons. Leptin-excited PMv-LepRb neurons are solely prescribed with 100 nM of leptin and 100 nM of SOCS3. Upon leptin binding, the baseline of resting membrane potential is hyperpolarized. However, with the activation of *I*_TRPC_ channels and inhibition of *I*_KCa_ channels by prolactin, these neurons show slightly increased spiking frequency and elongated bursting duration (Fig. 5C). Since PMv-LepRb neurons response to leptin (pSTAT3-ir) remains unchanged during pregnancy (45), we predict DAT expression and the proportion of leptin-excited (75%) *vs* leptin-inhibited PMv neurons (25%) will remain unchanged.

The increased spiking frequency and elongated bursting duration of leptin-inhibited PMv-LepRb neurons (Fig. 5C) induce glutamate release into AgRP presynaptic terminals. Due to regulation of action potentials by feed-forward and presynaptic inputs (73), AgRP neurons spiking frequency is significantly increased, and the depolarized duration is also significantly prolonged (Fig. 6C). The actions of leptin-excited PMv-LepRb neurons onto Kiss1 neurons are not changed by increased prolactin due to the low coexpression of PrlR and LepRb/non-DAT (leptin-depolarized) neurons. On the other hand, AgRP/GABA innervation of Kiss1 neurons is amplified by leptin-inhibited PMv-LepRb/DAT neurons, decreasing their activity and restraining the reproductive axis (Fig. 6F). This is an expected outcome in adaptive physiology to negative energy balance and high prolactin levels. Thus, we postulate that inhibition of Kiss1 neurons by AgRP/GABA inputs may function as an important counter-regulatory element for the leptin-activated PMv-LepRb neurons in conditions of reproductive restraint.

In this study, we showed that the PMv LepRb/DAT neurons are hyperpolarized by leptin and have proposed a neuronal circuitry in which they act to inhibit the neuroendocrine reproductive axis. Because DAT mRNA expression is decreased in adult females, the use of the DAT-Cre mouse model for experimental activation or inhibition of PMv LepRb/DAT neurons is not appropriate due to the striking decrease in DAT mRNA expression in adult mice and the difference in the colocalization rate between the transcript and Cre-activity. To circumvent the lack of experimental tools, we modelled the actions of PMv DAT and non-DAT LepRb neurons using a physiological and well-described condition of reproductive restraint. Late pregnancy and lactation are characterized by a temporary suppression of the reproductive axis and high levels of prolactin, necessary for milk production and maternal behavior (15, 26, 27, 41-44). High prolactin induces central leptin resistance decreasing hypothalamic response to the increased levels of leptin for the necessary accumulation of fat mass (15, 28). In the PMv, however, leptin’s effect seems to be preserved (45) and, together with the stimulatory effects of prolactin, it may contribute to the inhibition of the reproductive neuroendocrine axis in conditions of high energy demands. Experimental studies are necessary to test our model.

## Methods

### Experimental design

All procedures were carried out in accordance with the National Research Council Guide for the Care and Use of Laboratory Animals, and protocols were approved by the University of Michigan IACUC (PRO00008712). Adult DAT-Cre tdTom males (n=14) and females (n=6) were used to determine sex differences in the number of neurons expressing the reporter gene. Adult wild type (WT) male (n=3) and female (n=4) mice were used to study sex differences in DAT (*Slc6a3*) gene expression. WT mice were also used to determine developmental differences in *Slc6a3* gene expression, i.e. prepubertal (P19, n=7) vs adult (P60-70, n=5) diestrous female mice, and to assess the effects of estradiol (E2) on *Slc6a3* gene expression, i.e. ovariectomized (OVX, n=5), OVX + E2 (n=5) and diestrous females (n=4). Diestrous WT female mice (n=3) were used to assess DAT, LepR and PrlR mRNA co-expression by fluorescent *in situ* hybridization. Details on mice lines, cohorts and housing can be found in SI Appendix.

### Ovariectomy and estradiol replacement

Females underwent bilateral ovariectomy. OVX females received steroid replacement via a Silastic capsule containing E2 (1 μg, OVX+E2) or oil (OVX) subcutaneously at the time of surgery. OVX females were perfused 7-14 days following surgery, while OVX+E2 females were perfused 2 days following E2 replacement. Details on surgeries and controls can be found in SI Appendix.

### In situ hybridization (ISH)

Coronal fixed-frozen sections (30-μm) from perfused mice, at 120µm resolution, were used for radioactive ISH, as previously described (74, 75), using an ^35^S-UTP labelled dopamine transporter (DAT) riboprobe (750bp). The following primers were used (exon 10-15 of the *Slc6a3* gene): Forward (5’ ACGTCTTGATCACTGGGCTTGTCGATGAGTT 3’) and reverse (5’ GCATGGATTGGGTGTGAACAGTC 3’). Details on tissue preparation, ISH, imaging and quantification can be found in SI Appendix.

### Leptin treatment

DAT-Cre tdTomato mice fasted overnight were injected with ip. leptin (2.5 mg/kg, purchased from Dr. Parlow) (n = 3 females; n = 5 males). Forty-five minutes following leptin injection, mice were perfused, and tissue sections 150-µm apart were processed for pSTAT3 immunohistochemistry as described (24, 74). Tissue was incubated in primary rabbit anti-pSTAT3 (1:1,000; Cell Signaling) for 48 h at 4°C and detected using diaminobenzidine (DAB) and nickel ammonium sulfate (Sigma) as chromogens. Details of the method, quantification and imaging in SI Appendix.

### Electrophysiological recordings

Coronal sections from hypothalamic blocks (250 µM) from adult DAT-Cre tdTom male mice were prepared and the data analyzed as previously described (33). The pipette solution was modified to include an intracellular dye (Alexa Fluor 488) for whole-cell recording: 120 mM K-gluconate, 10 mM KCl, 10 mM HEPES, 5 mM EGTA, 1 mM CaCl_2_, 1 mM MgCl_2_, 2 mM (Mg)-ATP, and 0.03 mM AlexaFluor 488 hydrazide dye, pH 7.3. Whole-cell patch-clamp recordings were performed on tomato-positive (red fluorescence) neurons anatomically restricted to the PMv. In current-clamp mode, tdTom neurons were recorded under zero current injection (I = 0) in whole-cell patch-clamp configuration. The recording electrodes had resistances of 5-7 MΩ when filled with the K-gluconate internal solution. The membrane potential values were compensated to account for the junction potential (−8 mV). The resting membrane potential (RMP) was monitored for at least 10-15 minutes (baseline period) before leptin was administered to the bath. Solutions containing leptin (100 nM) were typically perfused for 5 min after the baseline period. Details on section preparation, recording and analysis can be found in SI Appendix.

### Tracing PMv-DAT neuronal projections

DAT-Cre mice received unilateral stereotaxic injections of an adenoassociated virus (AAV) expressing synaptophysin-mCherry fusion protein (25 nL) in the PMv. Three weeks after surgery mice were perfused and brains were harvested for histology. Fixed coronal 30-µm hypothalamic brain sections were processed for immunofluorescence. Primary antibodies used were rabbit polyclonal anti-AgRP (1:5,000, Phoenix Pharmaceuticals H-003-57) and rat monoclonal anti-mCherry 16D7 (1:5000, Invitrogen M11217). Details in SI Appendix.

### Fluorescent ISH

ISH was performed on fresh frozen 16-µm thick cryostat sections at 128-µm resolution (8-series). The ISH was performed following the RNAscope (ACDbio, RNAscope Multiplex Fluorescent Reagent Kit v2) protocol for fresh frozen sections, using Protease III. Probes used were: Mm-Slc6a3-C1 (#315441), Mm-Prlr-C2 (#430791-C2) and Mm-Lepr-C3 (#402731-C3, labeling all *LepR* isoforms). Details on the procedure, imaging and quantification are described in SI Appendix.

### Mathematical model and numerical methods

Each of the 4 neuronal types in the proposed model is treated as a lumped cell with conductance-based electrophysiology. The integrated dynamical system with 19 states is solved for the biological duration of 160 seconds by Matlab stiff ODE solver. Detailed modeling for ionic conductance of each neuronal type is formulated in SI Appendix and SI Appendix Tables S1, S2, S3, S4.

### Data Sharing

The manuscript contains all information necessary for replication of the data, including additional details in Supplemental Information. If additional information is required, including the code used in the model simulation, data will be readily available for editors, reviewers, and readers upon request.

## Acknowledgements

We thank Dr. Michael Roberts for the critical review of the manuscript, Dr. Yun-Hee Choi for the design of the DAT riboprobe, and Susan Allen for expert technical assistance. Dr. A. F. Parlow, Harbor-UCLA Medical Center, Torrance, California, USA; through the National Hormone and Peptide Program, for the leptin. This work was supported by the National Institutes of Health (R01-HD-069702 to CFE, CSM and NB), CNPq (Brazilian National Council for Scientific and Technological Development fellowship to BCB), the Coordenação de Aperfeiçoamento de Pessoal de Nível Superior – Brasil (CAPES) – Finance Code 001 (MAS) and by the São Paulo Research Foundation [FAPESP-Brazil, grants number: 13/07908-8 (RF), 15/20198-5 (TTZ).

