## Supplementary material for "Reproductive neuronal circuitry in adaptive changes of energy balance": SI Appendix

### Supplementary Information

#### Methods

***Animals.*** Adult (3-6 months of age) male and female mice expressing Cre-recombinase under the DAT promoter (JAX®; Stock 006660), a ROSA26 stop-floxed tdTomato reporter mouse line (R26 tdTomato, JAX®; Stock 007914), and wild type C57B6/J mice were used for experiments (JAX® mice, stock # 000664). Mice were housed in an Association for Assessment and Accreditation of Laboratory Animal Care (AAALAC) accredited facility at the University of Michigan Medical School under a 12 h light/dark cycle, temperature-controlled environment (21-23 °C), and were fed on a low-phytoestrogen diet (Envigo 2016 diet). All procedures were carried out in accordance with the National Research Council Guide for the Care and Use of Laboratory Animals, and protocols were approved by the University of Michigan IACUC (PRO00008712).

***Experimental design.*** Several cohorts of wild type (WT) male and female mice were used to assess DAT gene expression in the PMv. A cohort of adult DAT-Cre tdTom males (n=14) and females (n=6) were used to determine sex differences in the number of neurons expressing the reporter gene. One cohort of adult WT male (n=3) and female (n=4) mice was used to study sex differences in DAT (*Slc6a3*) gene expression. Another cohort of WT mice were used to determine developmental differences in DAT (*Slc6a3*) gene expression, ie. prepubertal (P19, n=7) and adult (P60-70, n=5) diestrous female mice. A third cohort of WT mice was used to assess the effects of estradiol (E2) on DAT (*Slc6a3*) gene expression, ie. ovariectomized (OVX, n=5), OVX + E2 (n=5) and diestrous females (n=4). An additional cohort of diestrous female mice (n=3) was used to assess DAT, LepR and PrlR mRNA co-expression by fluorescent *in situ* hybridization.

***Tissue collection and histology.*** Mice were anesthetized with isoflurane and transcardially perfused with diethylpyrocarbonate (DEPC)-treated saline followed by 10% neutral buffered formalin. After perfusion, brains were collected and postfixed in 20 % sucrose-10 % formalin for 2-4 h. Brains were then switched to a 20 % sucrose-DEPC/PBS solution overnight. Coronal 30-µm thick sections of the forebrain were acquired using a freezing microtome (Leica SM 2010R). Brain slices were divided in 4 series, were placed in RNAse-free cyroprotectant and were stored at -20 °C. For RNAscope, mice were anesthetized with isofluorane and immediately decapitated.

Brains were snap frozen. Coronal 16-μm thick sections of the forebrain were acquired using a cryostat (Leica CM 3050S) and mounted onto Superfrost Plus slides (Fisher). Sections were divided in 8 series and stored at -80 °C until further processing.

***Ovariectomy and estradiol replacement.*** Females were deeply anesthetized with isoflurane and underwent bilateral ovariectomy. A group of OVX females received steroid replacement implanted via a Silastic capsule containing E2 (1 μg, OVX+E2) or oil (OVX) subcutaneously at the time of surgery. OVX females were perfused, as described above 7-14 days following surgery, while OVX+E2 females were perfused 2 days following E2 replacement. Uterus size was used as control for the treatment. Only OVX mice with uterine weight below 80 mg and OVX+E2 mice with uterine weight above 100 mg were used. Both groups were perfused in the morning to avoid time of day effects of E2 feedback.

***Leptin treatment.*** One group of DAT-Cre tdTomato mice were fasted overnight and injected with ip. leptin (2.5 mg/kg, purchased from Dr. Parlow through the National Hormone and Peptide Program) (n = 3 females; n = 5 males). Forty-five minutes following leptin injection, mice were perfused, and tissue was processed as described above. Brain sections were pretreated with 30% $H_2O_2$ to block endogenous peroxidase activity, 0.3% glycine and 0.03% sodium dodecyl sulphate prior to primary antibody incubation. Tissue was blocked in 3% normal donkey serum, 0.25% Triton in PBS for 1 hour and incubated in primary rabbit anti-pSTAT3 (Cell Signaling, 1:1000) for 48 h at 4°C. Primary antibody was detected by avidin-biotin peroxidase method (Biotin-SP-conjugated donkey anti-rat; 1:1000; Jackson ImmunoResearch and ABC Kit, Vector Laboratories) using diaminobenzidine (DAB; Sigma) as chromogen. Quantification of tdTomato positive neurons was made by an observer unaware of the experimental groups. Dual labeled tdTomato and pSTAT3 immunoreactive cells were counted in each individual channel and colocalization was considered when pSTAT3 immunoreactivity was clearly nuclear in tdTomato positive cells. Two consecutive sections on one side of the brain with a representative image of the PMv were counted. No correction for double counting was performed because sections are 120 μm apart.

***Electrophysiological recordings.*** Hypothalamic slices were prepared and the data analyzed as previously described (8). Briefly, mice were decapitated, and the entire brain was removed. After removal, the brains were immediately submerged in ice-cold, carbogen-saturated (95% $O_2$ and

5% $CO_2$) ACSF (126 mM NaCl, 2.8 mM KCl, 26 mM $NaHCO_3$, 1.25 mM $NaH_2PO_4$, 1.2 mM $MgSO_4$, 5 mM glucose and 2.5 mM $CaCl_2$). Coronal sections from hypothalamic blocks (250 µM) were cut with a Leica VT1000S vibratome and incubated in oxygenated ACSF at room temperature for at least 1 hour before the recordings. The slices were transferred to the recording chamber and allowed to equilibrate for 10–20 min. The slices were bathed in oxygenated ACSF (32°C) at a flow rate of ~2 mL/min. The pipette solution was modified to include an intracellular dye (Alexa Fluor 488) for whole-cell recording: 120 mM K-gluconate, 10 mM KCl, 10 mM HEPES, 5 mM EGTA, 1 mM $CaCl_2$, 1 mM $MgCl_2$, 2 mM (Mg)-ATP, and 0.03 mM AlexaFluor 488 hydrazide dye, pH 7.3. Whole-cell patch-clamp recordings were performed on tomato-positive (red fluorescence) neurons anatomically restricted to the PMv of male DAT-Cre tdTom mice (8-12 weeks). Epifluorescence was briefly used to target the fluorescent cells, at which time the light source was switched to infrared differential interference contrast imaging to obtain the whole-cell recording (Leica DM6000 FS equipped with a fixed stage and a Leica DFC360 FX high-speed monochrome fluorescence digital camera). The electrophysiological signals were recorded using an Axopatch 700B amplifier (Molecular Devices), low-pass filtered at 2–5 kHz, and analyzed offline on a PC with the pCLAMP program (Molecular Devices). The recording electrodes had resistances of 5-7 MΩ when filled with the K-gluconate internal solution. The input resistance was assessed by measuring the voltage deflection at the end of the response to a hyperpolarizing rectangular current pulse (500 ms of -10 to -50 pA). The membrane potential values were compensated to account for the junction potential (-8 mV). The resting membrane potential (RMP) was monitored for at least 10-15 minutes (baseline period) before leptin (purchased through the National Hormone and Peptide Program) were administered to the bath. Solutions containing leptin (100 nM) were typically perfused for 5 min after the baseline period.

***Tracing PMv-DAT neuronal projections.*** DAT-Cre mice were anesthetized with isoflurane (2%, inhalation), were placed in a stereotaxic apparatus (Knopf) and the skull was exposed. Intracranial injection coordinates to the PMv were measured from the rostral rhinal vein [anterior-posterior: -0.54], the exposed superior sagittal sinus [medial-lateral: ± 0.04], and from the top of the dura-mater [dorsal-ventral -0.53]. Adeno-associated virus (AAV) containing the synapstophysin-mCherry fusion protein (25 nL) was unilaterally delivered into the PMv using a picospritzer attached to a glass pipette that was inserted into the brain. All mice were allowed to recover for 21 days prior to perfusion.

***In situ hybridization (ISH).*** The dopamine transporter (DAT) riboprobe was produced from mouse hypothalamic cDNA. The following primers were used to amplify a 754 base-pair sequence in the gene encoding for the DAT (exon 10-15 of the *Slc6a3* gene): Forward (5' ACGTCTTGATCACTGGGCTTGTCGATGAGTT 3') and reverse (5' GCATGGATTGGGTGTGAACAGTC 3'). Sequences for T7 and T3 promoters were added to primer sequences. Single labeled *in situ* hybridization was performed on 30-µm thick fixed brain sections mounted onto Superfrost Plus slides from adult WT male mice (P60-70), prepubertal female mice (P19), adult intact diestrous (P60-70), OVX, and OVX+E2 mice. Sections were subjected to a 10 min microwave sodium citrate (pH 6.0) pre-treatment and hybridized with an $^{35}S$-labeled DAT probe, as previously described (54, 55). Slides were dipped in autoradiographic emulsion (Kodak), dried for 3 h and stored in light-protected boxes at 4 °C for one week. Slides were developed in D-19 developer, dehydrated in ethanol, cleared in xylene, and coverslipped with DPX (Electron Microscopy Sciences). Images were acquired using digital camera on an AxioImager M2 microscope (Zeiss). *In situ* hybridization signals were quantified using integrated optical density (IOD) and ImageJ software (NIH). IODs were calculated from a constant area and IOD of tissue background was subtracted.

***Fluorescent ISH.*** For colocalization analysis, ISH was performed on fresh frozen 16-µm thick cryostat sections using an adaptation of the RNAscope (ACDbio, RNAscope Multiplex Fluorescent Reagent Kit v2) protocol for fresh frozen sections (n=4 diestous females). Briefly, sections were rinsed in PBS for 5 min, fixed in 10 % NBF for 2h at 4 °C, dehydrated through serial ethanol, cleared in xylene for 15 min and rehydrated on serial ethanol. Target retrieval was performed by baking the slides for 10 min in boiling sodium citrate buffer (0.01M, pH 6.0) and treating with 0.03 % sodium dodecyl sulphate for 3 min. Slides were dehydrated as before and air-dried for 20 min. ISH was performed using the RNAscope Protease III (ACDBio). Sections were incubated with Mm-Slc6a3-C1 (#315441), Mm-Prlr-C2 (#430791-C2) and Mm-Lepr-C3 (#402731-C3, labeling all *LepR* isoforms) RNAscope probes for 2 h at 40 °C using the HybEZ Humidifying System (ACDBio). After all incubation steps, slides were incubated in DAPI solution (320850) for 30 s at room temperature, and coverslipped using ProLong Gold Antifade Mountant (ThermoFisher Scientific). Quantification of mRNA coexpression within cells was performed on PMv images acquired at 40 × on an AxioImager M2 microscope (M2) and analysed using Zen 3.1 software (Zeiss). Based on observed background outside of the area of interest, a threshold for a minimum number of puncta per cell was used to consider a cell positive for expression of a specific gene. Confocal images were acquired on a Nikon A1 confocal microscope using set acquisition conditions.

***Immunofluorescence.*** Fixed 30-µm brain sections were rinsed in PBS, blocked with 3% normal donkey serum in PBS + 0.25% Triton-X 100 and incubated overnight at 4°C with primary antibodies. Primary antibody was visualized using secondary fluorescent antibodies. Antibodies used were rabbit polyclonal anti-AgRP (1:5,000, Phoenix Pharmaceuticals H-003-57), rat monoclonal anti-mCherry 16D7 (1:5000, Invitrogen M11217); and secondary donkey anti-rabbit AlexaFluor 488 and donkey anti-rat AlexaFluor 594 (1:500; Invitrogen).

***Mathematical model and numerical methods.*** The proposed PMv energy balancing circuitry is composed of four neuronal types. Each type is treated as a lumped cell with conductance-based electrophysiology. Detailed modeling for ionic conductance of each neuronal type is formulated in the next section. The relay from upstream neuron of leptin-excited and leptin-inhibited PMv-LepRb neurons to Kiss1 and AgRP neurons are by neurotransmitter/neuropeptide release dependent on the presynaptic excitation and mass-action kinetics of postsynaptic receptor gating. The integrated dynamical system with 19 states is solved for the biological duration of 160 seconds by Matlab stiff ODE solver.

#### *Conductance-based electrophysiology*

### A. Leptin-excited PMv-LepRb neurons

$$I_{\text{ionic}} = I_{\text{TRPC}} + I_{\text{CaL}} + I_{\text{K}} + I_{\text{KCa}} + I_{\text{leak}} \quad [1]$$

$$\frac{dV}{dt} = -I_{\text{ionic}}/C_{\text{m}} \quad [2]$$

#### 1. TRPC channel

$$I_{\text{NSC,Na}} = P_{\text{NSC,Na}}^{\text{max}} \cdot A_{\text{m}} \cdot \frac{F}{eK_{\text{B}}T} \cdot V \cdot \frac{[\text{Na}^+]_{\text{in}} - [\text{Na}^+]_{\text{out}} e^{-\frac{V}{eKbT}}}{1 - e^{-\frac{V}{eKbT}}} \times 10^6 \quad [3]$$

$$I_{\text{NSC,Ca}} = P_{\text{NSC,Ca}}^{\text{max}} \cdot A_{\text{m}} \cdot \frac{4F}{eK_{\text{B}}T} \cdot V \cdot \frac{[\text{Ca}^{2+}]_{\text{in}} - [\text{Ca}^{2+}]_{\text{out}} e^{-\frac{2V}{eKbT}}}{1 - e^{-\frac{2V}{eKbT}}} \times 10^6 \quad [4]$$

$$\alpha = \frac{\frac{[\text{leptin}]}{K_{\text{leptin}}}}{1 + \frac{[\text{leptin}]}{K_{\text{leptin}}} + \frac{[\text{SOCS3}]}{K_{\text{SOCS3}}}} \quad [5]$$

$$I_{\text{TRPC}} = \alpha(I_{\text{NSC,Na}} + I_{\text{NSC,Ca}}) \quad [6]$$

#### 2. L-type calcium channel

$$I_{\text{CaL}} = g_{\text{CaL}} \cdot m_\infty \cdot (V - 120) \quad [7]$$

$$m_\infty = (1 + \tanh\left(\frac{V + 1.2}{18}\right))/2 \quad [8]$$

#### 3. Voltage-sensitive potassium channel

$$I_{\text{K}} = g_{\text{K}} \cdot n \cdot (V + 84) \quad [9]$$

$$n_\infty = (1 + \tanh\left(\frac{V - 1.2}{17.4}\right))/2 \quad [10]$$

$$\tau_n = 1/\cosh\left(\frac{V - 12}{34.8}\right) \quad [11]$$

$$\frac{dn}{dt} = \phi \frac{n_\infty - n}{\tau_n} \quad [12]$$

#### 4. Calcium-activated potassium channel

$$I_{\text{KCa}} = g_{\text{SK}} \cdot \frac{[\text{Ca}^{2+}]_{\text{in}}}{1 + [\text{Ca}^{2+}]_{\text{in}}} \cdot (V + 84) \quad [13]$$

#### 5. Leakage

$$I_{\text{leak}} = g_{\text{leak}} \cdot (V + 60) \quad [14]$$

#### 6. Calcium handling

$$\frac{d[Ca^{2+}]_{\text{in}}}{dt} = \epsilon(-\mu(I_{\text{CaL}} + \alpha I_{\text{NSC,Ca}}) - [\text{Ca}^{2+}]_{\text{in}}) \quad [15]$$

**Table S1. Leptin-excited PMv-LepRb neurons**

| Parameter | Definition | Value | Reference |
|---|---|---|---|
| $C_m$ | Plasma membrane capacitance | 1.1 pF | (*1*) |
| $A_m$ | Plasma membrane surface area | 14.0 x $10^{-6}$ $cm^2$ | (*1*) |
| $T$ | Temperature | 300 K | |
| $eK_BT$ | electron energy in voltage | 24.8 mV | |
| $K_{leptin}$ | Half saturation for leptin | 1 $\mu$M | (*2*) |
| $K_{SOCS3}$ | Half saturation for SOCS3 | 1 $\mu$M | (*3*) |
| [leptin] | Concentration of leptin | 100 nM | (*4*) |
| [SOCS3] | Concentration of SOCS3 | 100 nM | (*4, 5*) |
| $P_{NSC,Na}^{max}$ | Maximum permeability of $Na^+$ in NSC channel | 3.214 x $10^{-7}$ cm/s | (*1*) |
| $P_{NSC,Ca}^{max}$ | Maximum permeability of $Ca^{2+}$ in NSC channel | 1.445 x $10^{-7}$ cm/s | (*1*) |
| $[Na^+]_{in}$ | Intracellular $Na^+$ concentration | 8.3 mM | (*6*) |
| $[Na^+]_{out}$ | Extracellular $Na^+$ concentration | 140.0 mM | (*6*) |
| $[Ca^{2+}]_{out}$ | Extracellular $Ca^{2+}$ concentration | 2.0 mM | (*6*) |
| $g_{CaL}$ | maximum conductance of L-type $Ca^{2+}$ channel | 4 nS | (*7*) |
| $g_K$ | maximum conductance of voltage-sensitive $K^+$ channel | 8 nS | (*7*) |
| $g_{SK}$ | maximum conductance of SK channel | 2 nS | (*7*) |
| $g_{leak}$ | maximum conductance of leakage | 2 nS | (*7*) |
| $\epsilon$ | Balancing factor between incoming calcium and uptake | 1.667 x $10^{-4}$ | (*7*) |
| $\mu$ | Conversion factor from incoming current to lumped intracellular concentration | 0.02 | (*7*) |
| $\phi$ | Scaling factor in potassium channel gating kinetics | 4.6 | (*7*) |
| $V_p$ | Presynaptic membrane voltage threshold for glutamergic release to Kiss1 | -33.0 mV | (*8*) |
| $g_{release}^{max}$ | Presynaptic glutamergic maximum release to Kiss1(75 % population) | 0.3 mM | (*8*) |

### B. Leptin-inhibited PMv-LepRb neurons

$$I_{\text{ionic}} = I_{\text{Na}} + I_{\text{TRPC}} + I_{\text{CaL}} + I_{\text{CaT}} + I_{\text{K}} + I_{\text{h}} + I_{\text{KCa}} + I_{\text{KATP}} + I_{\text{leak}} \quad [16]$$

$$\frac{dV}{dt} = -I_{\text{ionic}}/C_{\text{m}} \quad [17]$$

#### 1. Voltage-sensitive sodium channel

$$I_{\text{Na}} = g_{\text{Na}} \cdot m_\infty^3 \cdot (1-n) \cdot (V-50) \quad [18]$$

$$m_\infty = (1 + \tanh\left(\frac{V+1.2}{18}\right))/2 \quad [19]$$

$$n_\infty = (1 + \tanh\left(\frac{V-1.2}{17.4}\right))/2 \quad [20]$$

$$\tau_n = 1/\cosh\left(\frac{V-12}{34.8}\right) \quad [21]$$

$$\frac{dn}{dt} = \phi\frac{n_\infty - n}{\tau_n} \quad [22]$$

#### 2. TRPC channel

$$I_{\text{NSC,Na}} = P_{\text{NSC,Na}}^{\text{max}} \cdot A_{\text{m}} \cdot \frac{F}{\text{eK}_{\text{B}}T} \cdot V \cdot \frac{[\text{Na}^+]_{\text{in}} - [\text{Na}^+]_{\text{out}}\text{e}^{-\frac{V}{\text{eKb}T}}}{1-\text{e}^{-\frac{V}{\text{eKb}T}}} \times 10^6 \quad [23]$$

$$I_{\text{NSC,Ca}} = P_{\text{NSC,Ca}}^{\text{max}} \cdot A_{\text{m}} \cdot \frac{4F}{\text{eK}_{\text{B}}T} \cdot V \cdot \frac{[\text{Ca}^{2+}]_{\text{in}} - [\text{Ca}^{2+}]_{\text{out}}\text{e}^{-\frac{2V}{\text{eKb}T}}}{1-\text{e}^{-\frac{2V}{\text{eKb}T}}} \times 10^6 \quad [24]$$

$$\alpha = \frac{\frac{[\text{prolactin}]}{K_{\text{prolactin}}}}{1+\frac{[\text{prolactin}]}{K_{\text{prolactin}}}+\frac{[\text{SOCS2}]}{K_{\text{SOCS2}}}} \quad [25]$$

$$I_{\text{TRPC}} = \alpha(I_{\text{NSC,Na}} + I_{\text{NSC,Ca}}) \quad [26]$$

#### 3. L-type calcium channel

$$I_{\text{CaL}} = \beta_{\text{DAT}} \cdot g_{\text{CaL}} \cdot m_\infty \cdot (V-120) \quad [27]$$

#### 4. T-type calcium channel

$$I_{\text{CaT}} = g_{\text{CaT}} \cdot M_{\text{CaT},\infty}^2 \cdot H_{\text{CaT},\infty} \cdot (V-120) \quad [28]$$

$$M_{\text{CaT},\infty} = \frac{1}{1+e^{-(V+56.1)/10}} \quad [29]$$

$$H_{\text{CaT},\infty} = \frac{1}{1+e^{-(V+86.4)/4.7}} \quad [30]$$

#### 5. Voltage-sensitive potassium channel

$$I_{\text{K}} = g_{\text{K}} \cdot n \cdot (V+84) \quad [31]$$

#### 6. Hyperpolarization-activated channel

$$I_{\text{h}} = \beta_{\text{DAT}} \cdot g_{\text{h}} \cdot q \cdot (V+34.4) \quad [32]$$

$$q_\infty = \frac{1}{1+e^{(V+90.3)/9.67}} \quad [33]$$

$$\tau_q = \frac{1}{0.00062\cdot(e^{\frac{V+68}{-22}}+e^{\frac{V+68}{7.14}})} \quad [34]$$

$$\frac{dq}{dt} = \frac{q_\infty - q}{\tau_q} \quad [35]$$

### 7. Calcium-activated potassium channel

$$I_{\mathrm{KCa}} = (1 - \alpha) \cdot g_{\mathrm{SK}} \cdot \frac{[\mathrm{Ca}^{2+}]_{\mathrm{in}}}{1+[\mathrm{Ca}^{2+}]_{\mathrm{in}}} \cdot (V + 84) \quad [36]$$

### 8. ATP-dependent potassium channel

$$I_{\mathrm{KATP}} = \gamma \cdot g_{\mathrm{kATP}} \cdot p0_{\mathrm{ATP}} \cdot (V + 84) \quad [37]$$

$$p0_{\mathrm{ATP}} = \frac{0.08 \cdot (1+\frac{2[\mathrm{MgADP}]}{0.01}+0.89 \cdot \left(\frac{[\mathrm{MgADP}]}{0.01}\right)^2}{\left(1+\frac{[\mathrm{MgADP}]}{0.01}\right)^2 (1+\frac{0.45 \cdot [\mathrm{MgADP}]}{0.026}+\frac{[\mathrm{ATP}]}{0.05})} \quad [38]$$

$$\gamma = \frac{\frac{[\mathrm{leptin}]}{K_{\mathrm{leptin}}}}{1+\frac{[\mathrm{leptin}]}{K_{\mathrm{leptin}}}+\frac{[\mathrm{SOCS3}]}{K_{\mathrm{SOCS3}}}} \quad [39]$$

### 9. Leakage

$$I_{\mathrm{leak}} = g_{\mathrm{leak}} \cdot (V + 25.05) \quad [40]$$

### 10. Calcium handling

$$\frac{d[\mathrm{Ca}^{2+}]_{\mathrm{in}}}{dt} = \epsilon(-\mu\left(I_{\mathrm{CaL}} + I_{\mathrm{CaT}} + \alpha I_{\mathrm{NSC,Ca}}\right) - [\mathrm{Ca}^{2+}]_{\mathrm{in}}) \quad [41]$$

**Table S2. Parameter values in leptin-inhibited PMv-LepRb neurons**

| Parameter | Definition | Value | Reference |
|---|---|---|---|
| $C_\mathrm{m}$ | Plasma membrane capacitance | 1.1 pF | (*1*) |
| $K_\mathrm{prolactin}$ | Half saturation for prolactin | 50 $\mu$M | (*9*) |
| $K_\mathrm{leptin}$ | Half saturation for leptin | 1 $\mu$M | (*2*) |
| $K_\mathrm{SOCS2}$ | Half saturation for SOCS2 | 5 $\mu$M | (*3*) |
| $K_\mathrm{SOCS3}$ | Half saturation for SOCS3 | 1 $\mu$M | (*3*) |
| [prolactin] | Concentration of prolactin | 5 $\mu$M | (*5, 10, 11*) |
| [leptin] | Concentration of leptin | 100nM | (*4*) |
| [SOCS2] | Concentration of SOCS2 | 1 $\mu$M | (*4, 5*) |
| [SOCS3] | Concentration of SOCS3 | 100nM | (*4, 5*) |
| $P_{\mathrm{NSC,Na}}^{\mathrm{max}}$ | Maximum permeability of $Na^+$ in NSC channel | 1.301 x $10^{-7}$ cm/s | (*1*) |
| $P_{\mathrm{NSC,Ca}}^{\mathrm{max}}$ | Maximum permeability of $Ca^{2+}$ in NSC channel | 5.845 x $10^{-6}$ cm/s | (*1*) |
| $g_\mathrm{K}$ | maximum conductance of voltage-sensitive $K^+$ channel | 10 nS | (*7*) |
| $g_\mathrm{Na}$ | maximum conductance of $Na^+$ channel | 28 nS | (*7*) |
| $g_\mathrm{CaL}$ | maximum conductance of L-type $Ca^{2+}$ channel | 4 nS | (*7*) |
| $g_\mathrm{CaT}$ | maximum conductance of T-type $Ca^{2+}$ channel | 0.4 mS | (*7*) |
| $g_\mathrm{h}$ | maximum conductance of hyperpolarization-activated channel | 912 nS | (*7*) |
| $g_\mathrm{kATP}$ | maximum conductance of ATP-dependent $K^+$ channel | 8.741 $\mu S$ | (*12*) |
| $g_\mathrm{SK}$ | maximum conductance of SK channel | 1.187 nS | (*7*) |
| $g_\mathrm{leak}$ | maximum conductance of leakage | 2 nS | (*7*) |
| [MgADP] | Concentration of MgADP | 0.127 mM | (*12*) |
| [ATP] | Concentration of ATP | 2.65 mM | (*12*) |
| $\beta_{\mathrm{DAT}}$ | Normalized DAT expression | 1.2 | |
| $V_\mathrm{p}$ | Presynaptic membrane voltage threshold for glutamergic release to AgRP | -5.0 mV | (*8*) |
| $\epsilon$ | Balancing factor between incoming calcium and uptake | 1.667 x $10^{-4}$ | (*7*) |
| $\mu$ | Conversion factor from incoming current to lumped intracellular concentration | 0.02 | (*7*) |
| $\phi$ | Scaling factor in potassium channel gating kinetics | 4.6 | (*7*) |
| $g_{\mathrm{release}}^{\mathrm{max}}$ | Presynaptic glutamergic maximum release to AgRP (25 % population) | 0.1 mM | (*8*) |

### C. Kisspeptin neurons

$$I_{\text{ionic}} = I_{\text{Na}} + I_{\text{CaL}} + I_{\text{CaT}} + I_{\text{K}} + I_{\text{KCa}} + I_{\text{h}} + I_{\text{leak}} \quad [42]$$

$$\frac{dV}{dt} = -(I_{\text{ionic}} + I_{\text{NMDA}} + I_{\text{GABA}})/C_{\text{m}} \quad [43]$$

#### 1. Voltage-sensitive sodium channel

$$I_{\text{Na}} = g_{\text{NaP}} \cdot m_{\text{Na},\infty} \cdot h \cdot (V - 50) \quad [44]$$

$$m_{\text{Na},\infty} = \frac{1}{1 - e^{(V+40)/6}} \quad [45]$$

$$h_{\infty} = \frac{1}{1 + e^{(V+48)/6}} \quad [46]$$

$$\tau_h = \bar{\tau}_h / \cosh\left(\frac{V+48}{12}\right) \quad [47]$$

$$\frac{dh}{dt} = \frac{h_{\infty} - h}{\tau_h} \quad [48]$$

#### 2. Voltage-sensitive L-type calcium channel

$$I_{\text{CaL}} = g_{\text{CaL}} \cdot m_{\infty} \cdot (V - 120) \quad [49]$$

$$m_{\infty} = (1 + \tanh\left(\frac{V+1.2}{18}\right))/2 \quad [50]$$

#### 3. Voltage-sensitive potassium channel

$$I_{\text{K}} = g_{\text{K}} \cdot n \cdot (V + 84) \quad [51]$$

$$n_{\infty} = (1 + \tanh\left(\frac{V-1.2}{17.4}\right))/2 \quad [52]$$

$$\tau_n = 1/\cosh\left(\frac{V-12}{34.8}\right) \quad [53]$$

$$\frac{dn}{dt} = \phi \frac{n_{\infty} - n}{\tau_n} \quad [54]$$

#### 4. Voltage-sensitive T-type calcium channel

$$I_{\text{CaT}} = g_{\text{CaT}} \cdot M_{\text{CaT},\infty}^2 \cdot H_{\text{CaT},\infty} \cdot (V - 120) \quad [55]$$

$$M_{\text{CaT},\infty} = \frac{1}{1 + e^{-(V+56.1)/10}} \quad [56]$$

$$H_{\text{CaT},\infty} = \frac{1}{1 + e^{-(V+86.4)/4.7}} \quad [57]$$

#### 5. Calcium-activated potassium channel

$$I_{\text{KCa}} = g_{\text{SK}} \cdot \frac{[\text{Ca}^{2+}]_{\text{in}}}{1 + [\text{Ca}^{2+}]_{\text{in}}} \cdot (V + 84) \quad [58]$$

#### 6. Hyperpolarization-activated channel

$$I_{\text{h}} = g_{\text{h}} \cdot q_{\text{h}} \cdot (V + 34.4) \,/\, 40 \quad [59]$$

$$q_{\text{h},\infty} = \frac{1}{1 + e^{(V+90.3)/9.67}} \quad [60]$$

$$\tau_{qh} = \frac{1}{0.00062 \cdot (e^{\frac{V+68}{-22}} + e^{\frac{V+68}{7.14}})} \quad [61]$$

$$\frac{dq_h}{dt} = \frac{q_{h,\infty} - q_h}{\tau_{qh}} \quad [62]$$

#### 7. Leakage

$$I_{\text{leak}} = g_{\text{leak}} \cdot (V + 60) \quad [63]$$

### 8. Synaptic currents

$$I_{\text{NMDA}} = g_{\text{NMDA}}^{\max} \cdot g_{\text{Kiss1,NMDA}} \cdot (V - V_{\text{NMDA}}) \quad [64]$$

$$I_{\text{GABA}} = g_{\text{GABA}}^{\max} \cdot g_{\text{Kiss1,GABA}} \cdot (V - V_{\text{GABA}}) \quad [65]$$

### 9. Calcium handling

$$\frac{d[\text{Ca}^{2+}]_{\text{in}}}{dt} = \epsilon(-\mu(I_{\text{CaL}} + I_{\text{CaT}}) - [\text{Ca}^{2+}]_{\text{in}}) \quad [66]$$

**Table S3. Parameter values in Kisspeptin neurons**

| Parameter | Definition | Value | Reference |
|---|---|---|---|
| $C_{\text{m}}$ | Plasma membrane capacitance | 1.1 pF | (*1*) |
| $g_{\text{NaP}}$ | maximum conductance of persistent $Na^+$ channel | 0.5 nS | (*7*) |
| $g_{\text{K}}$ | maximum conductance of voltage-sensitive $K^+$ channel | 8 nS | (*7*) |
| $g_{\text{CaL}}$ | maximum conductance of L-type $Ca^{2+}$ channel | 4 nS | (*7*) |
| $g_{\text{CaT}}$ | maximum conductance of T-type $Ca^{2+}$ channel | 4 nS | (*7*) |
| $g_{\text{h}}$ | maximum conductance of hyperpolarization-activated channel | 400 nS | (*7*) |
| $g_{\text{kATP}}$ | maximum conductance of ATP-dependent $K^+$ channel | 2.31 n*S* | (*12*) |
| $g_{\text{SK}}$ | maximum conductance of SK channel | 0.75 nS | (*7*) |
| $g_{\text{leak}}$ | maximum conductance of leakage | 2 nS | (*7*) |
| $V_{\text{NMDA}}$ | reversal potential for the current from NMDA receptor | 30.0 mV | (*13*) |
| $V_{\text{GABA}}$ | reversal potential for the current from GABA receptor | -80.0 mV | (*8*) |
| $g_{\text{NMDA}}^{\max}$ | maximum conductance of NMDA channel | 2.0 nS | (*13*) |
| $g_{\text{GABA}}^{\max}$ | maximum conductance of GABA channel | 2.0 nS | (*8*) |
| $\bar{\tau}_h$ | Relaxation time for inactivating variable | 10 s | (*7*) |
| $\epsilon$ | Balancing factor between incoming calcium and uptake | 0.002 | (*7*) |
| $\mu$ | Conversion factor from incoming current to lumped intracellular concentration | 0.019 | (*7*) |
| $\phi$ | Scaling factor in potassium channel gating kinetics | 4.6 | (*7*) |
| $r_{\text{open}}^{\text{NMDA}}$ | NMDA receptor opening rate | 1.25 /mM/ms | (*8*) |
| $r_{\text{closed}}^{\text{NMDA}}$ | NMDA receptor closing rate | 0.045 /ms | (*8*) |
| $r_{\text{open}}^{\text{GABA}}$ | GABA receptor opening rate | 2.5 /mM/ms | (*8*) |
| $r_{\text{closed}}^{\text{GABA}}$ | GABA receptor closing rate | 0.09 /ms | (*8*) |

### D. AgRP neurons

$$I_{\mathrm{ionic}} = I_{\mathrm{Na}} + I_{\mathrm{CaL}} + I_{\mathrm{K}} + I_{\mathrm{Kir}} + I_{\mathrm{KCa}} + I_{\mathrm{kATP}} + I_{\mathrm{leak}} \quad [67]$$

$$\frac{dV}{dt} = -(I_{\mathrm{ionic}} + I_{\mathrm{NMDA}})/C_{\mathrm{m}} \quad [68]$$

#### 1. Voltage-sensitive sodium channel

$$I_{\mathrm{Na}} = g_{\mathrm{Na}} \cdot m_\infty \cdot (1-n) \cdot (V-50) \quad [69]$$

$$m_\infty = (1 + \tanh\left(\frac{V+1.2}{18}\right))/2 \quad [70]$$

$$n_\infty = (1 + \tanh\left(\frac{V-1.2}{17.4}\right))/2 \quad [71]$$

$$\tau_n = 1/\cosh\left(\frac{V-12}{34.8}\right) \quad [72]$$

$$\frac{dn}{dt} = \phi\frac{n_\infty - n}{\tau_n} \quad [73]$$

#### 2. Voltage-sensitive potassium channel

$$I_{\mathrm{K}} = g_{\mathrm{K}} \cdot n \cdot (V+84) \quad [74]$$

#### 3. Inward-rectifier potassium channel

$$I_{\mathrm{Kir}} = g_{\mathrm{Kir}} \cdot n_{\mathrm{Kir},\infty} \cdot (V+84) \quad [75]$$

$$n_{\mathrm{Kir},\infty} = \frac{0.8}{1+e^{(V+80)/12}} + 0.2 \quad [76]$$

#### 4. ATP-sensitive potassium channel

$$I_{\mathrm{KATP}} = \alpha \cdot g_{\mathrm{kATP}} \cdot p0_{\mathrm{ATP}} \cdot (V+84) \quad [77]$$

$$p0_{\mathrm{ATP}} = \frac{0.08\cdot(1+\frac{2[\mathrm{MgADP}]}{0.01}+0.89\cdot\left(\frac{[\mathrm{MgADP}]}{0.01}\right)^2}{\left(1+\frac{[\mathrm{MgADP}]}{0.01}\right)^2(1+\frac{0.45\cdot[\mathrm{MgADP}]}{0.026}+\frac{[\mathrm{ATP}]}{0.05})} \quad [78]$$

$$\alpha = \frac{\frac{[\mathrm{leptin}]}{K_{\mathrm{leptin}}}}{1+\frac{[\mathrm{leptin}]}{K_{\mathrm{leptin}}}+\frac{[\mathrm{SOCS3}]}{K_{\mathrm{SOCS3}}}} \quad [79]$$

#### 5. Voltage-sensitive L-type calcium channel

$$I_{\mathrm{CaL}} = g_{\mathrm{CaL}} \cdot m_\infty \cdot (V-120) \quad [80]$$

#### 6. Calcium-dependent potassium channel

$$I_{\mathrm{KCa}} = g_{\mathrm{SK}} \cdot \frac{[\mathrm{Ca}^{2+}]_{\mathrm{in}}}{1+[\mathrm{Ca}^{2+}]_{\mathrm{in}}} \cdot (V+84) \quad [81]$$

#### 7. Leakage

$$I_{\mathrm{leak}} = g_{\mathrm{leak}} \cdot (V+60) \quad [82]$$

#### 8. Synaptic current

$$I_{\mathrm{NMDA}} = g_{\mathrm{NMDA}}^{\max} \cdot g_{\mathrm{AgRP,NMDA}} \cdot (V - V_{\mathrm{NMDA}}) \quad [83]$$

#### 9. Calcium handling

$$\frac{d[\mathrm{Ca}^{2+}]_{\mathrm{in}}}{dt} = \epsilon(-\mu I_{\mathrm{CaL}} - [\mathrm{Ca}^{2+}]_{\mathrm{in}}) \quad [84]$$

**Table S4. Parameter values in AgRP neurons**

| Parameter | Definition | Value | Reference |
|---|---|---|---|
| $C_m$ | Plasma membrane capacitance | 1.1 pF | (*1*) |
| $K_{leptin}$ | Half saturation for leptin | 1 $\mu$M | (*2*) |
| $K_{SOCS3}$ | Half saturation for SOCS3 | 1 $\mu$M | (*3*) |
| [leptin] | Concentration of leptin | 100 nM | (*4*) |
| $g_{Na}$ | maximum conductance of $Na^+$ channel | 26.6 nS | (*7*) |
| $g_K$ | maximum conductance of voltage-sensitive $K^+$ channel | 8 nS | (*7*) |
| $g_{Kir}$ | maximum conductance of inward rectifier $K^+$ channel | 0.02 nS | (*7*) |
| $g_{CaL}$ | maximum conductance of L-type $Ca^{2+}$ channel | 4 nS | (*7*) |
| $g_{kATP}$ | maximum conductance of ATP-dependent $K^+$ channel | 0.924 $\mu$S | (*12*) |
| $g_{SK}$ | maximum conductance of SK channel | 0.9375 nS | (*7*) |
| $g_{leak}$ | maximum conductance of leakage | 2 nS | (*7*) |
| [MgADP] | Concentration of MgADP | 0.127 mM | (*12*) |
| [ATP] | Concentration of ATP | 2.65 mM | (*12*) |
| $V_{NMDA}$ | reversal potential for the current from NMDA receptor | -5.0 mV | (*13*) |
| $g_{NMDA}^{max}$ | maximum conductance of NMDA channel | 0.1 $\mu$S | (*13*) |
| $\epsilon$ | Balancing factor between incoming calcium and uptake | 0.001 | (*7*) |
| $\mu$ | Conversion factor from incoming current to lumped intracellular concentration | 0.01 | (*7*) |
| $\phi$ | Scaling factor in potassium channel gating kinetics | 5.52 | (*7*) |
| $r_{open}^{NMDA}$ | NMDA receptor opening rate | 1.25 /mM/ms | (*8*) |
| $r_{closed}^{NMDA}$ | NMDA receptor closing rate | 0.045 /ms | (*8*) |
| $V_p$ | Presynaptic membrane voltage threshold for GABAergic release to Kiss1 | -20.0 mV | (*8*) |
| $g_{release}^{max}$ | Presynaptic GABAergic maximum release to Kiss1 | 1.0 mM | (*8*) |
